# Genetic markers of ADHD-related variations in intracranial volume

**DOI:** 10.1101/184192

**Authors:** Marieke Klein, Raymond K. Walters, Ditte Demontis, Jason L. Stein, Derrek P. Hibar, Hieab H. Adams, Janita Bralten, Nina Roth Mota, Russell Schachar, Edmund Sonuga-Barke, Manuel Mattheisen, Benjamin M. Neale, Paul M. Thompson, Sarah E. Medland, Anders D. Børglum, Stephen V. Faraone, Alejandro Arias-Vasquez, Barbara Franke

## Abstract

Attention-Deficit/Hyperactivity Disorder (ADHD) is a common and highly heritable neurodevelopmental disorder with a complex pathophysiology, where genetic risk is hypothesized to be mediated by alterations in structure and function of diverse brain networks. We tested one aspect of this hypothesis by investigating the genetic overlap between ADHD (n=55,374) and (mainly subcortical) brain volumes (n=11,221-24,704), using the largest publicly available studies. At the level of common variant genetic architecture, we discovered a significant negative genetic correlation between ADHD and intracranial volume (ICV). Meta-analysis of individual variants found significant loci associated with both ADHD risk and ICV; additional loci were identified for ADHD and amygdala, caudate nucleus, and putamen volumes. Gene-set analysis in the ADHD-ICV meta-analytic data showed significant association with variation in neurite outgrowth-related genes. In summary, our results suggest new hypotheses about biological mechanisms involved in ADHD etiology and highlight the need to study additional brain parameters.

## INTRODUCTION

Attention-Deficit/Hyperactivity Disorder (ADHD) is a common and highly heritable neurodevelopmental disorder with a complex and presumably heterogeneous pathophysiology. Pathways towards disease are hypothesized to be mediated by alterations in diverse brain networks^1^. A recent neuroimaging mega-analysis reported subtle but consistent differences in the volumes of subcortical brain regions and intracranial volume (ICV) in ADHD, across diverse cohorts worldwide: compared to healthy controls, patients with ADHD showed decreased volumes of the nucleus accumbens, amygdala, caudate nucleus, hippocampus, putamen, and ICV^2^. How such alterations contribute to the disease phenotype, is still poorly understood. However, brain volume alterations are also present, on average, in unaffected relatives of patients with ADHD^3,4^, and both ADHD and brain volumes have high heritability (60-70%^5-7^ and 70-90%^8^, respectively). This suggests that genetic variants underlying ADHD pathophysiology may also influence brain volume variation. Recently, the first genome-wide significant loci for ADHD were identified and a SNP-based heritability of 20.16% has been reported^9^. The current study aimed to determine, if common genetic variants are shared between ADHD risk and subcortical brain volumes or ICV.

Genome-wide association studies (GWAS) have identified ten genome-wide significant loci associated with hippocampal volume^8,10-13^, eight loci with ICV^8,10,14,15^, four loci with putamen volume^8^, and one locus with caudate volume^8^. These variants explain only a small fraction of the heritability of the brain structure volumes^8,10-12,14^. Recently, Franke and colleagues reported a battery of statistical tools to comprehensively examine genetic overlap between brain volumes and risk for brain disease at the genome-wide level and at the level of individual risk variants, using schizophrenia as an example^16^. Here, we applied a similar set of methods to identify and dissect genetic sharing between ADHD and brain volumes implicated in ADHD based on the latest mega-analysis^2^. We used the largest available data sets, i.e., the GWAS meta-analysis (GWAS-MA) of ADHD from the Psychiatric Genomics Consortium (PGC) and The Lundbeck Foundation Initiative for Integrative Psychiatric Research (iPSYCH)^9^ and the GWAS-MAs of brain volumes from the ENIGMA Consortium (Enhancing NeuroImaging Genetics through Meta-Analysis)^8,13,17^ and the Neurology Working Group of the Cohorts for Heart and Aging Research in Genomic Epidemiology (CHARGE) Consortium^18,19^.

## RESULTS

We analyzed GWAS-MA data for ADHD (20,183 cases and 35,191 controls) and six MRI-based measures of brain volumes (11,221–24,704 individuals; subcortical brain volumes were corrected for ICV). Detailed information on the numbers of individuals and genetic variants of these different cohorts are given in **Supplementary Table 1**. These data were used in a comprehensive set of analyses of common variant genetic sharing between ADHD and brain volumetric measures evaluating potential overlap at global and single variant levels.

### Comparison of common variant genetic architectures

*Linkage disequilibrium score regression*. The SNP-based heritability estimates for the MRI measures were consistent with prior reports^13,16,19^ and ranged from 13.32% (nucleus accumbens) to 28.15% (putamen; Table 1). The amygdala mean *χ*^2^ was too small to allow a valid analysis. In the analysis of the genetic correlations, we observed a significant negative genetic correlation between ADHD and ICV (rg=-0.227, P=0.00015). All other correlations were non-significant (Table 1; **Supplementary Table 2** shows the results when using constrained intercepts).

**Table 1.** SNP heritability analyses for MRI brain volumes and genetic correlation with ADHD^*^.

| Brain region | N | Heritability | SE | Genetic correlation with ADHD | SE | Z | P |
| --- | --- | --- | --- | --- | --- | --- | --- |
| Nucleus accumbens | 11,709 | 0.1332 | 0.0518 | 0.005558 | 0.09487 | 0.05858 | 0.9533 |
| Caudate nucleus | 11,772 | 0.2456 | 0.0455 | -0.06426 | 0.07321 | -0.8778 | 0.3801 |
| Hippocampus<sup>#</sup> | 24,704 | 0.1418 | 0.0286 | -0.02354 | 0.06902 | -0.3411 | 0.733 |
| Intracranial volume<sup>#</sup> | 24,024 | 0.2318 | 0.0325 | -0.2266 | 0.05989 | -3.784 | **0.0001543** |
| Putamen | 11,646 | 0.2815 | 0.056 | 0.006433 | 0.07077 | 0.09089 | 0.9276 |
| Hippocampus ENIGMA only | 11,665 | 0.1363 | 0.0488 | -0.04202 | 0.09965 | -0.4217 | 0.6733 |
| Intracranial volume ENIGMA only | 11,221 | 0.1745 | 0.0461 | -0.2348 | 0.09452 | -2.485 | 0.01296 |

\*Amygdala mean $\chi^2$ was too small to allow a valid analysis (N=11,757). <sup>#</sup>Using GWAS-MA summary statistics from the meta-analysis of ENIGMA and CHARGE cohorts. Heritability and genetic correlation were estimated by using free intercepts. In accordance with the number of tests performed, we set a Bonferroni-corrected significance level at $P=0.05/6=0.0083$ . P-values in bold are significant after Bonferroni correction.

*SNP effect concordance analysis*. We used SNP Effect Concordance Analysis (SECA)^20^ to determine the extent and directionality of genetic overlap between ADHD and each brain volume. We found significant evidence of global pleiotropy for variants affecting ADHD risk for volumes of four subcortical brain regions and ICV (ADHD pleiotropy only reached nominal significance for the nucleus accumbens, Table 2 and **Supplementary Fig. 1**). The discordant SNP effects for ADHD and ICV were significant, i.e. variants increasing the risk for ADHD were associated with decreased ICV (PICV<0.001; Table 2 and **Supplementary Fig. 2**). Evidence for concordant SNP-effects reached significance for ADHD and nucleus accumbens (P_accumbens_=0.002) and caudate nucleus (P_caudate_=0.004, Table 2 and **Supplementary Fig. 2**); analyses for other structures did not reach study-wide significance (Table 2).

**Table 2.** Results of pleiotropy and concordance test of SNP Effect Concordance Analysis. Brain volume GWAS-MA was conditioned on ADHD GWAS-MA.

| Brain volume | P<sub>pleiotropy</sub> | CI<sub>pleiotropy</sub> | P<sub>concordance</sub> | CI<sub>concordance</sub> | Direction of SNP effects |
| --- | --- | --- | --- | --- | --- |
| Nucleus accumbens | 0.034 | 0.0244-0.0471 | **0.002** | 0.000548-0.00726 | concordant |
| Amygdala | **<0.001** | 5.12x10<sup>-5</sup>-0.00564 | 0.006 | 0.00275-0.013 | discordant |
| Caudate nucleus | **<0.001** | 5.12x10<sup>-5</sup>-0.00564 | **0.004** | 0.00156-0.0102 | concordant |
| Hippocampus<sup>#</sup> | **0.002** | 0.000548-0.00726 | 1 | 0.996-1 | / |
| Intracranial volume<sup>#</sup> | **<0.001** | 5.12x10<sup>-5</sup>-0.00564 | **<0.001** | 5.12x10<sup>-5</sup>-0.00564 | discordant |
| Putamen | **<0.001** | 5.12x10<sup>-5</sup>-0.00564 | 0.01 | 0.00544-0.0183 | concordant |
| Hippocampus ENIGMA only | 0.005 | 0.00214-0.0116 | 1 | 0.996-1 | / |
| Intracranial volume ENIGMA only | **<0.001** | 5.12x10<sup>-5</sup>-0.00564 | **<0.001** | 5.12x10<sup>-5</sup>-0.00564 | discordant |

P-values and confidence intervals (CI) were obtained based on 1,000 permutations. <sup>#</sup>Using GWAS-MA summary statistics from the meta-analysis of ENIGMA and CHARGE cohorts. In accordance with the number of tests performed, we set a Bonferroni-corrected significance level at $P=0.05/(2*6)=0.00416$ . P-values in bold are significant after Bonferroni correction.

*Sign tests*. Based on the phenotypic observation that patients with ADHD have, on average, smaller brain volumes compared to healthy controls^2^, we had expected discordant rather than concordant SNP effects. As both discordant and concordant effects were observed in the SECA among the brain volumes, we subsequently used sign tests to specifically determine directionality of genetic overlap between ADHD and each brain volume for the top-associations per trait. In this, we zoomed in further on the most strongly associated and LD-independent SNPs and compared the signs of the regression coefficients of those top-associations per trait across ADHD and the MRI volumetric measures. None of the sign tests showed a consistent direction of discordance, after correction for multiple testing (**Supplementary Table 3**).

### Analyses at the single genetic variant level

*Weighted SNP meta-analyses*. Based on the findings of both concordant and discordant links between ADHD and the brain volume SNPs, we performed a genome-wide search for specific genetic loci associated with both ADHD and each of the brain traits. For this, we used a weighted SNP metaanalysis design allowing the combination of findings from GWAS of binary and quantitative variables^9^. This enabled us to specifically look for concordant effects at the level of single genetic variants; there is currently no suitable method to study discordant effects. The weighted GWAS-MA for ADHD and ICV, comprising 79,398 individuals from independent cohorts, identified two loci of interest, one on chromosome 15 (*SEMA6D*) and one on chromosome 16 (intergenic). Those loci were genome-wide significant in the cross-phenotype meta-analysis and/or had a cross-phenotype p-value, which was improved by at least one order of magnitude, and a cross-phenotype z-score, which (at least) equaled the ones observed in the GWAS-MAs for the individual phenotypes (Table 3 and Figures 1 and 2). Four additional LD-independent genetic loci passed the study-wide threshold for genome-wide significance of P=8.33x10^−9^, but those were related to a single phenotype and did not meet the criteria for cross-disorder relevance (Figure 1).

**Figure 1:**
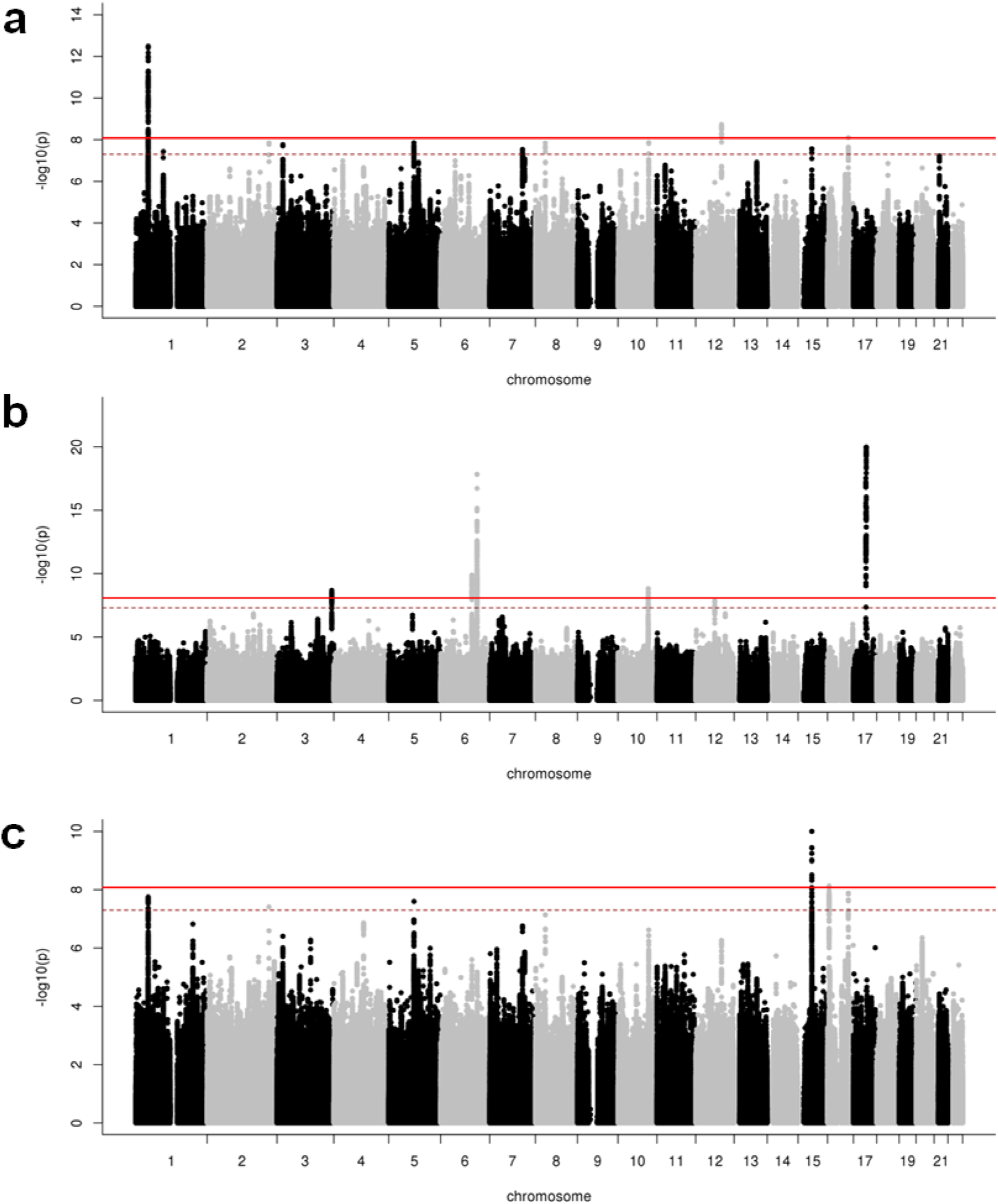
Common genetic variants associated with ADHD, ICV and ADHD+ICV. Shown here are Manhattan plots, in which every point represents a single genetic variant plotted according to its genomics position (x-axis) and its –log10(P) for association with the respective trait (y-axis). The solid bright red line represents the study-wide genome-wide significance of P<8.33x10^−9^, and the dashed dark red line represents the genome-wide significance of P<5x10^−8^. (a) PGC+iPSYCH ADHD GWAS-MA. (b) ENIGMA+CHARGE ICV GWAS-MA. (c) ADHD+ICV weighted GWAS-MA.

We also performed weighted GWAS-MAs for ADHD and four subcortical brain structures. For amygdala volume, a naïve sample size-weighted meta-analysis was performed, as no genetic correlation with ADHD had been estimated. Manhattan plots for these analyses are in **Supplementary Figures 3-7**; the six novel and/or improved LD-independent genome-wide significant loci observed in these analyses are summarized in Table 3. Among those, the *SEMA6D* locus was significantly associated with ADHD and putamen volume (P=3.62x10^−9^; Table 3, Figure 2, and **Supplementary Fig. 7**).

**Table 3:** Results of (weighted) meta-analyses of ADHD and brain volumes showing novel or improved independent genome-wide significant loci.

| Brain trait | Chr | Pos | Gene | SNP | A1 | Zscore | P-value | Direction ADHD / brain | Zscore<sub>ADHD</sub> | Zscore<sub>brain</sub> |
| --- | --- | --- | --- | --- | --- | --- | --- | --- | --- | --- |
| Intracranial volume | 15 | 47769424 | *SEMA6D* | rs281320<sup>a</sup> | T | -6.4677 | 9.95x10<sup>-11</sup> | -/- | -5.5487 | -3.33 |
| Intracranial volume | 16 | 5829204 | intergenic | rs13332522<sup>b</sup> | C | 5.7810 | 7.43x10<sup>-9</sup> | +/+ | 4.6162 | 3.508 |
| Amygdala | 3 | 20725016 | intergenic | rs56135409<sup>a</sup> | A | -6.0105 | 1.85x10<sup>-9</sup> | -/- | -5.3525 | -2.7482 |
| Caudate nucleus | 5 | 87847273 | *LINC00461* | rs12653396<sup>b</sup> | A | 5.3826 | 1.46 x10<sup>-9</sup> | +/+ | 5.3777 | 2.9499 |
| Putamen | 15 | 47754027 | *SEMA6D* | rs281323<sup>a</sup> | A | 5.9005 | 3.62 x10<sup>-9</sup> | +/+ | 5.4775 | 2.2114 |
| Putamen | 2 | 184483159 | intergenic | rs114181198<sup>b</sup> | A | 5.1308 | 4.28 x10<sup>-9</sup> | +/+ | 5.0859 | 3.2596 |

Results for six genome-wide significant index SNPs identified in the GWAS meta-analyses of ADHD+brain volumes. The allele (A1) for the effect of the Z-score is given. Direction gives information about the direction of the association in the two samples, “+” indicates that A1 increases risk/volume and “-” indicates that A1 decreases risk/volume. The threshold for genome-wide significance was set at $P < 8.33 \times 10^{-9}$ ( $5 \times 10^{-8} / 6$ traits). <sup>a</sup>This locus improved by one order of magnitude after the meta-analysis. <sup>b</sup>This locus is a new genome-wide significant locus in the ADHD+brain volume meta-analysis.

**Figure 2:**
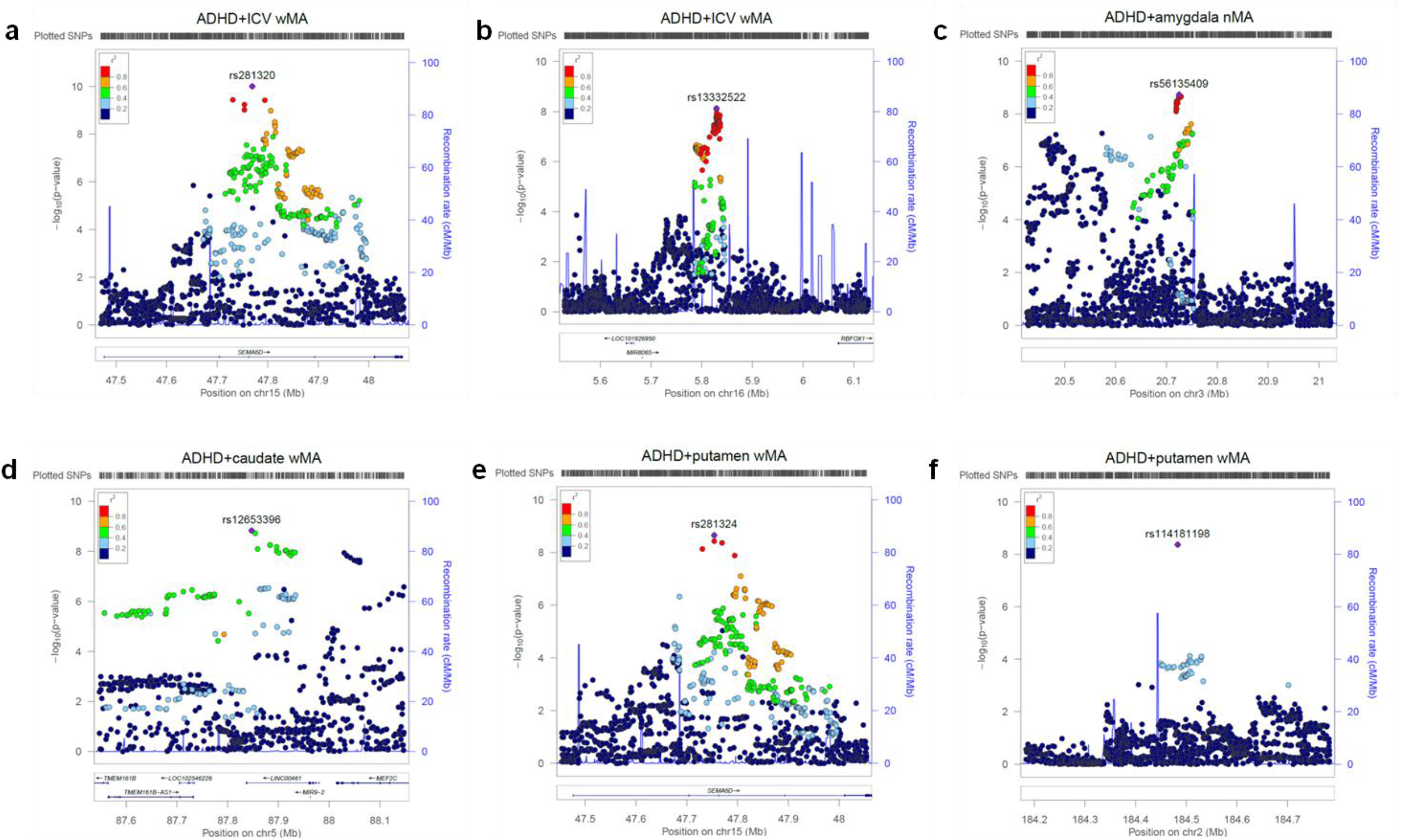
Regional association of genome-wide significant loci of ADHD and brain volume GWAS-MAs. For each panel (a-f), zoomed-in association plots (± 300 kb from the top SNP, indexed by purple dot) are shown. Plots are zoomed to highlight the genomic region that likely harbors the causal variant(s). Plots were made using the Locuszoom software^57^.

*Reciprocal lookup of genome-wide significant associations*. No significant associations were observed between the 14 previously identified genome-wide significant ADHD SNPs^9^ and brain volumes, (**Supplementary Table 4**). Conversely, among the 21 SNPs previously associated with the brain volumes^8,13,19^, two associations with ADHD survived correction for multiple testing (**Supplementary Table 5**; P<0.00238). A proxy variant (rs8756; r^2^=1; chr12:66359752, located in exon 5 of *HMGA2*) of the ICV-associated variant rs138074335 (chr12:66374247, intergenic and upstream of *HMGA2*; increasing ICV) was associated with increased risk for ADHD (OR=1.042, P=0.00186). A second variant, rs2195243 (chr12:102922986, intergenic and upstream of *IGF1*), was associated with decreased ICV and increased risk for ADHD (OR=1.0565, P=0.00127).

### Expression quantitative loci and brain gene expression

The recently published ADHD GWAS-MA paper described biological annotations implementing the co-localized genes for some of the genome-wide significant loci^9^. Among those, it was shown that many SNPs in the *SEMA6D* locus on chromosome 15, which we found overlapping between ADHD risk and ICV, were strongly associated with the expression of *SEMA6D* in fibroblasts^21^. Indeed, repeating this analysis for the most strongly associated variants identified by the (weighted) cross-phenotype SNP meta-analyses using the GTEx data^21^, we found that rs281320 on chromosome 15, the SNP with the strongest association in the ADHD+ICV GWAS-MA (P=9.95x10^−11^), to be a significant expression quantitative trait locus (eQTL) for *SEMA6D* in transformed fibroblast tissue (P=1.2x10^−20^; **Supplementary Table 6**). Likewise, rs281323 was also strongly associated with *SEMA6D* expression (P=1.2x10^−21^). The alternative alleles of both rs281320 and rs281323, which are associated with increased risk for ADHD and larger ICV, also increased *SEMA6D* expression (**Supplementary Fig. 9, a** and **b**). Additionally, rs12653396 was a significant eQTL for the *CTC-498M16.4* and *MEF2C* genes, in brain (P=2x10^−7^)^21^ and blood tissue (P=6.53x10^−7^)^22^, respectively, with the disease-associated A-allele also being associated with increased *MEF2C* expression (**Supplementary Table 6**). All other top-SNPs were not present in either of the two eQTL databases.

We determined mRNA expression for *SEMA6D* and *MEF2C*, the only protein-coding genes identified in the cross-phenotype GWAS-MAs, in six brain regions of the developing and adult human brain using data from the Human Brain Transcriptome Project^23^. Both genes were found to be globally expressed in the developing and adult human brain, with highest mRNA expression in prenatal periods (**Supplementary Fig. 8**).

### Neurite outgrowth gene-set analysis

Based on our finding that *SEMA6D* – a gene with a prominent role in neuronal migration and axonal path finding - is a key locus contributing to both ADHD risk and ICV, we investigated, whether genes involved in directed neurite outgrowth, more generally, have a role in ADHD–ICV genetic overlap. We found a significant association between a pre-defined gene-set of 45 neurite outgrowth genes^24^ and the meta-analytic data for ADHD+ICV using MAGMA (P_competitive_=0.00338). There were no significant associations with either ADHD or ICV, separately (**Supplementary Table 7**). As expected, we found a significant association of the neurite outgrowth gene-set with ADHD for the self-contained test (P_self-contained_=5.53x10^−9^). Using the ADHD+ICV GWAS-MA data, the most strongly associated individual gene among the set of neurite outgrowth genes was *CREB5* (P=0.000553), and a total of nine additional neurite outgrowth genes showed nominally significant associations (**Supplementary Table 8**).

## DISCUSSION

We found significant - though modest - genetic covariation between ADHD risk and brain volumes, both on global and single variant levels. On the global level, significant negative genetic correlation between ADHD and ICV was demonstrated. The direction of effect was supported by the SNP effect concordance analysis. For most of the subcortical brain volumes affected in ADHD, pleiotropic effects were also found. On the single variant level, meta-analyses found significant loci associated with both ADHD risk and brain volumes. We identified *SEMA6D* as a potential key-locus contributing to both ADHD risk and ICV, and gene-set analysis revealed a significant association of ADHD-ICV overlap with variation in previously ADHD-related neurite outgrowth genes.

In contrast to prior studies across different mental disorders, such as schizophrenia^15,18^, major depressive disorder^25^, and autism^19^, which provided no evidence for a sharing of genetic architecture and brain volumes, we found a modest but significant negative global genetic correlation between ADHD and ICV. This finding was highly consistent across approaches (i.e. LDSR and SECA; and reciprocal look-up for rs2195243) and in the expected direction, given the previous observation that patients with ADHD have smaller ICV relative to controls^2^. Recent studies also support a genetic correlation between ICV and adult height, Parkinson’s disease, and cognitive function in children and adults^19^. With ADHD being negatively genetically correlated with general cognitive function^26^ and human intelligence^27^, it remains to be tested, whether the same genes are involved.

We observed significant pleiotropy also between ADHD and amygdala, caudate nucleus, hippocampus, and putamen volumes. This global genetic covariation was substantiated by the local effects, which we observed in the weighted cross-phenotype meta-analyses. In addition to ICV, caudate nucleus, and putamen volumes variation also showed significant genetic concordance with ADHD. However, whereas the results for ICV were in line with our expectation (variants increasing risk for ADHD decrease brain volume; ADHD patients have overall smaller brain volume^2^), the concordant effects for ADHD and nucleus accumbens and caudate nucleus volume are counterintuitive (ADHD patients have smaller volumes for these structures^2^), suggesting a reverse or more complex pattern of causation.

We investigated genetic overlap on global and single variant levels. Combining both approaches is highly informative, as they answer different questions. Whereas on the global level, we are interested in the general genetic architecture underlying two phenotypes, the single variant level aims to identify specific loci that can inform us about shared biological mechanisms. In line with this, we performed a meta-analysis of ADHD and brain volumes even for those volumes that showed no detectable genetic correlation.

On the single variant level, we only had tools available to perform a meta-analysis by looking at concordant effects^9^, we therefore had to ignore locus-specific discordant effects. Still, the strongest association of single genetic markers was observed for ADHD and ICV, additional associations were identified for ADHD and the subcortical volumes. The weighted meta-analysis of ADHD and ICV found two potentially pleiotropic loci. One of those was *SEMA6D*, coding for the semaphorin 6D, a transmembrane molecule which plays a role in maintenance and remodeling of neuronal connections^28^. Animal studies have shown that it acts as a ligand for PlexinA1, which is involved in critical steps of neuronal development in the spinal cord^29^ and cardiac development^30,31^. This finding is consistent with prior research implicating neurite outgrowth dysregulation in ADHD^24,32^, and highlighting it as potential neural mediator of genetic risk for ADHD. This should be tested in future studies by applying imaging genetics mediation models^33^ or Mendelian randomization approaches^34^. We also found the *SEMA6D* locus in the cross-phenotype meta-analysis for ADHD and putamen volume, even though this volume had been corrected for ICV, suggesting that genetic variation in *SEMA6D* may influence specific brain regions to a different extent.

The current results raise a number of questions concerning the way that alterations in the brain mediate etiological risk pathways in ADHD. First, given the cross-sectional nature of the available data, we must be unbiased as to direction of causation. As long as we have not fully clarified the role of brain structure alterations in psychiatric disorders, i.e. whether those are cause or consequence of the disorders, we should keep an open mind regarding the direction of effects. Observed effects might not always be direct, but may involve reciprocal or transactional processes. Second, the degree of sharing was statistically modest. At first sight, this seems to be inconsistent with the general hypothesis that ADHD is a genetic-based brain disorder. However, there are a number of possible explanations for this modest amount of shared effects. First, we examined brain structure at a gross anatomical measure: atlas-based brain segmentations might be too coarse to identify subtler volumetric differences. In line with this, a surface-based analysis localized allelic effects of a SNP in *KTN1* to regions along the superior and lateral putamen^8^, suggesting that genetic variation does not affect structure across different regions of the brain in the same way. Additionally, cellular changes (e.g., altered spine density) may not manifest at such gross levels at all. Moreover, there is a limited link between structure and function at the level of the MRI-derived volumes, so other neuroimaging phenotypes (e.g., structural and functional connectivity measures) could be more informative. As pointed out previously^16^, the limited SNP-heritability of the (subcortical) brain volumes further challenges the identification of genetic overlap. Non-significance of some findings might also be related to sample size. However, based on the comparison of the LDSR results for ENIGMA-only and for ENIGMA+CHARGE, we can conclude that sample size affected the standard errors and therefore the significance of this analysis, but not the point estimates. On the other hand, the cross-phenotype meta-analysis results were mainly driven by the association signal of the larger GWAS-MA, namely the ADHD data set, although several loci improved after combining the two phenotypes. However, even larger GWAS-MA data sets are becoming available for the (sub)cortical regional brain volumes^35^, so that the disorder and brain GWAS-MAs start to become equally well-powered. Finally, it is also possible that some of ADHD’s association with reduced brain volumes is driven by environmental effects, either independently or in interaction with genetic factors. Adverse prenatal and postnatal environmental exposures have been linked to both ADHD and general brain growth and development^1^. In line with this idea, monozygotic twins discordant for ADHD were found to have significant epigenetic differences among genes highly expressed in the developing cerebellum, striatum, and thalamus, in addition to genes functionally related to neurodevelopment and neurotransmitter regulation^36^.

The findings described here need to be interpreted in light of several strengths and limitations. Previous brain imaging genetics studies in ADHD mainly focused on single genetic variant investigations, as recently reviewed by Klein et al.^37^, and were hampered by limited sample sizes. The current study combined the largest data sets available to investigate the genetic overlap between ADHD and brain volumes. For this, a complementary battery of statistical methods and analyses was used, allowing the study of genetic covariation on both global and single variant levels and across the entire genome. Nevertheless, some limitations apply. Firstly, this study focused on a limited set of mainly subcortical MRI measures, and future work should be extended to cortical regions and white matter integrity^3,38,39^. Secondly, for the cross-phenotype GWAS-MA, we used a recently described weighted meta-analysis method^9^. However, we observed that with low and moderate genetically correlated phenotypes, the association signals did generally not improve over a naïve meta-analysis, performed without adding additional weights (data not shown). In addition, we could only investigate concordant SNP effects with this method. Novel methods for joint analysis of summary statistics from GWASs of different phenotypes^40^ and co-localization approaches^41^ are currently being developed and might help in the future to overcome these limitations and increase power. Thirdly, it is possible that this study underestimates genetic correlations, as we did not take into account the known role of rare variants in the genetic architecture of ADHD^42-44^. In line with the omnigenic model for complex traits^45^, and as the heritability of complex traits and diseases is spread broadly across the genome^46,47^, future studies investigating heritability and genetic correlation in low-minor allele frequency and low-LD regions may identify stronger relationships between ADHD and brain volumes.

This is the first study to show significant global and single variant level genetic correlations derived from polygenic overlap between ADHD and brain volumes. The subtle but significant genetic overlap between ADHD and variation in brain volumes is consistent with models implicating alterations in brain structure in ADHD-related genetic risk pathways and stimulate new hypotheses about neurobiological mechanisms involved in ADHD. However, the rather modest overlap reinforces the need to consider other structural and functional brain metrics and explore the role of environmental factors.

## ONLINE METHODS

### Ethics statement

The current study used summary statistics of GWAS meta-analyses (GWAS-MA) that had been approved by the local ethics committees and had the required informed consents, as described in the earlier publications^8,9,13,19^.

### Participant samples

#### ADHD Working Group of the PGC and the ADHD iPSYCH-SSI-Broad collaboration

ADHD GWAS-MA summary statistics data were acquired from the ADHD Working Group of the PGC and the ADHD iPSYCH-SSI-Broad collaboration (n=55,374^9^, https://www.med.unc.edu/pgc/results-and-downloads). Detailed quality control and imputation parameters are described in the original publication^9^. Briefly, genotype imputation was done using the bioinformatic pipeline “ricopili” and with the pre-phasing/imputation stepwise approach implemented in IMPUTE2/SHAPEIT using the haplotypes from the 1000 Genomes Project, phase 3, version 5 (1KGP3v5)^48^ data. Association analyses using the imputed marker dosages were performed separately for the 11 PGC samples and the 23 waves in iPSYCH by an additive logistic regression model using PLINK v1.9^49^, with the derived principal components included as covariates as described in the original publication^9^. Subsequently, meta-analysis, including summary statistics from GWASs of the 23 waves in iPSYCH and 11 PGC samples, was conducted using an inverse-weighted fixed effects model. In total, 20,183 cases and 35,191 controls were used for the original analysis (**Supplementary Table 1**). Only SNPs with imputation quality (INFO score) >0.8 and MAF >0.01 were included in the meta-analysis. PGC+iPSYCH ADHD GWAS-MA summary statistics data only included markers which were supported by an effective sample size greater than 70% (8,047,420 markers)^9^.

#### ENIGMA

In the GWAS-MAs on subcortical volumes those volumes had been adjusted for ICV to identify specific genetic contributions to individual volumes. Five subcortical volumes and ICV were selected for this study since significant volume reductions in patients with ADHD compared to controls were reported^2^. Access to the summary statistics of ENIGMA can be requested via their website (http://enigma.ini.usc.edu/download-enigma-gwas-results/). For the initial GWAS-MA analysis, MRI brain scans and genome-wide genotype data were available for 11,840 subjects from 22 cohorts. Genomic data were imputed to a reference panel (1000 Genomes, phase1, v3 (1KGP1v3)^50^) comprising only European samples and with monomorphic SNPs removed. Imputation was performed at each site using MaCH for phasing and minimac for imputation^51^. Only SNPs with an imputation score of RSQ >0.5 and minor allele counts >10 within each site were included. Tests of association were conducted separately for eight MRI volumetric phenotypes (nucleus accumbens, amygdala, caudate nucleus, hippocampus, pallidum, putamen, thalamus and ICV) with the following covariates in a multiple linear regression framework: age, age2, sex, four MDS components (to account for population structure) and ICV (for subcortical brain phenotypes). GWA statistics from each of the 22 sites were combined using a fixed-effect inverse variance-weighted meta-analysis as implemented in METAL^52^. Prior to all analyses, a cohort including ADHD cases (NeuroIMAGE cohort, n=154) was removed from the ENIGMA data.

#### CHARGE

We obtained genome-wide GWAS-MA summary statistics data on ICV and hippocampal volume from the CHARGE Consortium (n=12,803 and n=13,039, respectively^13,19^) and CHARGE summary statistics data had been requested by the principal investigator of the study described by Adams et al.^19^. GWAS of ICV and hippocampal volumes were performed for each site separately, controlling for age, sex, and, when applicable age^2^, population stratification variables, study site, and diagnosis (when applicable). Summary statistics, including effect estimates of the genetic variant with ICV or hippocampal volume under an additive model, were exchanged to perform a fixed-effects metaanalysis weighting for sample size in METAL^52^. After the final meta-analysis, variants were excluded if they were only available for fewer than 5,000 individuals.

### Removal of duplicated individuals

Subject overlap between all PGC ADHD and ENIGMA cohorts was evaluated using a checksum algorithm to ensure the robustness of our results, given that some analyses could be sensitive to the presence of duplicate individuals. For each individual, ten checksum numbers were created based on ten batches of 50 SNP genotypes and compared between individuals from both consortia. Based on these comparisons no subjects needed to be removed from the data sets. As no Danish cohort was included in the ENIGMA or CHARGE study, we assumed that there is no sample overlap between cohorts studying brain volume and iPSYCH.

### GWAS meta-analysis of ENIGMA and CHARGE data sets

To increase the sample size for the hippocampal volume and ICV data, summary statistics of GWAS-MA results from ENIGMA^8^ (after removal of ADHD cases) and CHARGE^13,19^ were combined using a fixed-effects sample size-weighted meta-analysis framework as implemented in METAL^52^. After the final meta-analysis, variants were excluded if they were only available for fewer than 5,000 individuals or a MAF ≤0.005. After filtering, the meta-analyses results included more than 9,145,464 markers.

### Linkage disequilibrium score regression (LDSR)

For LDSR, each GWAS-MA data set underwent additional filtering. Only markers overlapping with HapMap Project Phase 3 SNPs and passing the INFO score ≥0.9 and MAF ≥0.01 filters were included (where available). SNPs with missing values, duplicate rs-numbers, too low a sample size (where available SNPs with an effective sample size less than 0.67 times the 90th percentile of sample size were removed), or that were strand-ambiguous - as well as indels - were removed. As described in the original ADHD GWAS-MA paper^9^, for LDSR analysis the European only subset was used (n_cases_=19,099 and n_controls_=34,194), since LDSR requires linkage disequilibrium [LD] data from a sample of comparable ethnic background). For the ENIGMA amygdala results, the mean *χ*^2^ was too low (1.0) to reliably estimate SNP heritability using LDSR.

The analyses used a two-step procedure with the LD-scoring analysis package^53^. An unconstrained regression estimated regression intercepts for each pair of phenotypes. Since we adopted protocols to exclude sample overlap, we also performed the analyses with regression intercept for the genetic correlation analysis defined as zero (**Supplementary Table 2**). To compute p-values, standard errors were estimated using a block jackknife procedure.

### SNP effect concordance analysis (SECA)

#### Post-processing of genetic data

To statistically compare the ADHD and six brain volume GWAS-MAs, we used SNPs passing quality control and filtering rules (for ADHD GWAS-MA INFO ≥0.9 and MAF ≥0.01 and for ENIGMA and CHARGE GWAS-MA RSQ ≥0.5 and MAF ≥0.005) in all data sets. With these data, we performed a clumping procedure in PLINK^54^ to identify an independent SNP from every LD block across the genome. The clumping procedure was performed separately for each of the brain volume GWAS-MAs using a 500 kb window, with SNPs in LD (*r*^2^ >0.2) in the European reference samples from the 1KGP1v3^50^. The SNP with the lowest p-value within each LD block was selected as the index SNP representing that LD block and all other SNPs in the LD block were dropped from the analysis. The result after applying the clumping procedure was sets of independent SNPs representing the total variation explained across the genome conditioned on the significance in each brain volume GWAS-MA. For each of these sets of SNPs, we then determined the corresponding ADHD GWAS-MA test statistic for each independent, index SNP and used these data sets for the subsequent analyses.

#### Tests of pleiotropy and concordance

We used SNP Effect Concordance Analysis (SECA)^20^ to determine the extent and directionality of genetic overlap between ADHD and each brain volume. Within SECA we performed a global test of pleiotropy using a binomial test at 12 p-value levels: P ≤(0.01, 0.05, 0.1, 0.2, 0.3, 0.4, 0.5, 0.6, 0.7, 0.8, 0.9). For a given brain volume and ADHD paired set, we separately ordered SNPs based on their p-value for association with each trait. For each of the 12 p-value levels, we determined the total number of SNPs overlapping between the two traits at each p-value threshold and compared that number to the expected random overlap under the null hypothesis of no pleiotropy using a binomial test. In total, 144 comparisons were performed. We tallied the number of comparisons with evidence of overlap at a nominally significant level of P ≤0.05. To evaluate the global level of pleiotropy, we generated 1,000 permuted data sets for a given brain volume to ADHD comparison and determined, if the number of significance thresholds with genetic overlap was significantly greater than chance.

Similarly, we estimated concordance, the agreement in SNP effect directions across two traits. We determined whether or not there was a significant (P ≤0.05) positive or negative trend in the effect of the overlapping SNPs at each of the 12 p-value thresholds. This was done using a two-sided Fisher’s exact test. The direction of effect for each SNP was determined by the sign of the SNP regression coefficient (OR or beta value) from each meta-analysis. In the ADHD GWAS-MA, an odds ratio >1 for a SNP indicates that the A1 reference allele was associated with an increased risk of developing ADHD (an odds ratio <1 indicates a protective allele). A positive Beta value for a SNP in a brain volume GWAS-MA indicates that the A1 reference allele of that SNP is associated with an increase in brain volume (a negative Beta value indicates that the A1 reference allele of that SNP is associated with a reduction in brain volume). We estimated the global level of concordance between a given brain trait and ADHD by generating 1,000 permuted data sets, repeating the Fisher’s exact test procedure, and determined if the number of significant overlapping thresholds was significantly greater than chance (see Nyholt et al., 2014^20^ for details of the SECA analysis).

In total, we tested for pleiotropy and concordance between ADHD and six brain volumes. In accordance with the number of tests performed, we set a Bonferroni-corrected significance level at P=0.05/(2*6)=4.17x10^−3^.

### Independent genome-wide significant markers and loci

LD-independent markers associated at P <1x10^−5^ were defined using the clump flag in PLINK v1.9^49^. Clumping was used to group additional associated markers within a 0.5 Mb window surrounding the index SNP. Markers were grouped to the index SNPs if they were also associated (P <0.001) and were in LD with the index SNP (r^2^ >0.1). A genome-wide significant locus was defined as the physical region containing the identified LD independent index SNPs and their correlated variants (r^2^>0.8) with P <0.001. Associated loci within 250kb of each other were merged. All LD statistics were calculated using the 1KGP3v5^48^ reference haplotypes.

### SNP sign test in the top GWAS-MA findings

To investigate a potential accumulation of same- or opposite-direction effects of SNPs between ADHD and brain volumes, we counted the number of *opposite* direction effects (as expected from the imaging results in Hoogman et al.^2^) for the top findings from the ADHD data set in the different brain structure data sets. The ADHD GWAS-MA data were clumped to define independent loci for all variants with P<1x10^−5^ in the ADHD GWAS-MA using 1KGP3v5^48^ data on European ancestry populations as reference.

The proportion of variants with discordant direction of effect in the individual brain GWAS-MA was evaluated using a binomial test against a null hypothesis of 0.5 (i.e. the signs are discordant between the two analyses by chance). This test was done for loci passing p-value thresholds of P<5x10^−8^ (14 LD-independent genome-wide significant SNPs), P<1x10^−6^ (44 LD-independent SNPs), and P<1x10^−5^ (132 LD-independent SNPs) in the ADHD GWAS-MA.

### Weighted meta-analysis of ADHD and brain volume data sets

Independent of the results of the global overlap analyses, we also performed meta-analyses combining the results from the ADHD GWAS-MA with results from GWAS-MAs of brain volumes (amygdala, nucleus accumbens, caudate nucleus, hippocampus, putamen, and ICV). This was done using a modified sample size-based weighting method, integrating the binary ADHD trait (ADHD risk) with the continuous trait (brain volume traits), as described in Demontis et al.^9^. For the metaanalyses, modified sample size-based weights were derived to account for the respective heritability, genetic correlation, and measurement scale of the GWASs. The adjusted samples sizes reflect differences in power between the studies due to measurement scale and relative heritability that is not captured by sample size. Thereby, the contribution of the continuous phenotype’s GWAS to the meta-analysis is reduced based on imperfect correlation with the dichotomous phenotype of interest (in this case ADHD risk). The adjustments are computed based on the sample and population prevalence of the dichotomous phenotype, the estimated SNP heritability of the two phenotypes (liability scale for dichotomous phenotype), and the genetic correlation between the two phenotypes, as well as the average SNP LD score, and the number of SNPs. Heritability and genetic correlation values to compute these weights are computed using LD score regression^53^ as described before. For a comprehensive description of the method for meta-analysis of continuous and dichotomous phenotype and notes on the implementation please see the supplementary information of the original ADHD GWAS-MA publication^9^. For all brain volumes, we also performed naive meta-analyses given the low genetic correlations with ADHD risk observed. Correcting for meta-analyzing six brain phenotypes with ADHD, we set the threshold for genome-wide significance at P=5x10^−8^/6=8.33x10^−9^.

### Reciprocal lookup of significant GWAS-MA loci

Evidence for an effect of the independent ADHD-associated SNPs on brain volume was studied through a look-up of results in the ENIGMA(+CHARGE) GWAS-MAs. LD-independent loci with corresponding index SNPs were obtained by clumping summary statistics of the ADHD GWAS-MA^9^. Similarly, effects of the 21 independent SNPs significantly associated with brain volumes as reported in the original publications of the brain volume GWAS-MAs^8,13,19^ on ADHD risk were looked-up in the ADHD GWAS-MA data. If the index variant was not present in the other data set, a proxy variant was selected by using LDlink (https://analysistools.nci.nih.gov/LDlink/). Given the number of tests performed, we set a Bonferroni-corrected significance levels at P=0.05/(14*8)=0.000446 for the lookup of ADHD SNPs in brain volume GWAS-MAs and at P=0.05/21=0.002381 for brain volume SNPs in ADHD GWAS-MA data.

### Expression quantitative trait loci and brain gene expression

To assess potential functionality in (brain) tissues, we tested the identified risk variants (Table 3) for association with gene expression. Expression quantitative trait loci (eQTL) were examined using data from the GTEx portal (https://www.gtexportal.org/home/)^21^. In addition, blood eQTL data were queried using the Blood eQTL Browser (http://genenetwork.nl/bloodeqtlbrowser/)^22^.

We also investigated the spatio-temporal expression pattern in brain tissue for genes with significantly associated variants in the approaches described above (Table 3) using data from the Human Brain Transcriptome Project (http://hbatlas.org). We assessed messenger RNA (mRNA) expression trajectories in six regions of the developing and adult human brain. Spanning periods from embryonic development to late adulthood, this data set provides genome-wide exon-level transcriptome data generated using the Affymetrix GeneChip Human Exon 1.0 SS Arrays from over 1,340 tissue samples sampled from both hemispheres of *postmortem* human brains (n=57)^23^. Gene expression over the lifespan from the spatio-temporal atlas was graphed using custom R scripts^23^.

### Gene-based and gene-set analyses for ADHD+ICV GWAS-MA data

Based on our finding that *SEMA6D* is a key locus contributing to both ADHD risk and ICV – a loci involved in neuronal migration and axonal path finding – we investigated, whether neurite outgrowth-related genes in general have a role in ADHD–ICV genetic overlap. Genome-wide summary statistics of the (i) weighted meta-analysis for ADHD and ENIGMA+CHARGE ICV GWAS-MA, (ii) the ADHD GWAS-MA, and (iii) the ENIGMA+CHARGE GWAS-MA on ICV were used as input for gene-based and gene-set analyses. For the ADHD+ICV GWAS-MA, only SNPs shared between the ADHD and ICV data set were included. Statistical analyses used the Multi-marker Analysis of GenoMic Annotation (MAGMA) software package (version 1.05)^55^. Genome-wide SNP data from a reference panel 1KGP1v3^50^ was annotated to NCBI Build 37.3 gene locations using a symmetric 100 kb flanking window. The gene annotation file was used to map genome-wide SNP data from the different studies, to assign SNPs to genes followed by the calculation of gene-based p-values. For the gene-based analyses, single SNP p-values within a gene were transformed into a gene-statistic by taking the mean of the χ ^2^-statistic among the SNPs in each gene. To account for LD, the 1KGP1v3^50^ was used as a reference to estimate the LD between SNPs within (the vicinity of) the genes (http://ctglab.nl/software/MAGMA/ref_data/g1000_ceu.zip). Subsequent to the genome-wide gene-based analysis, we also tested, whether genes in the neurite-outgrowth gene-set (defined previously, n_genes_=45; ref. ^24^) were jointly associated with results of the weighted meta-analytic data of ADHD+ICV using self-contained and competitive testing^56^. Since self-contained tests do not take into account the overall level of association across the genome, gene size, and gene density, we were particularly interested in the competitive test for the current analysis. Subsequently, we tested whether the gene-set was associated with the two individual data sets as well. In this, the same procedure was followed for analysis of the ADHD GWAS-MA and ENIGMA+CHARGE ICV GWAS-MA summary statistics individually. *Post-hoc*, the individual genes in the set were investigated, by reviewing gene test-statistics of the weighted ADHD+ICV GWAS-MA results. Genes were considered gene-wide significant, if they reached the adjusted Bonferroni correction threshold (P=0.05/45=0.00111). Subsequently, we reviewed gene-based associations in the ADHD GWAS-MA and ENIGMA+CHARGE ICV GWAS-MA results separately.

URLs

http://enigma.ini.usc.edu/download-enigma-gwas-results/

https://www.med.unc.edu/pgc/results-and-downloads

https://github.com/bulik/ldsc

https://neurogenetics.qimrberghofer.edu.au/SECA/

https://analysistools.nci.nih.gov/LDlink/

https://www.gtexportal.org/home/

http://hbatlas.org

http://ctglab.nl/software/MAGMA/ref_data/g1000_ceu.zip

http://locuszoom.sph.umich.edu/

## DATA AVAILABILITY

The genome-wide summary statistics that support the findings of this study are available at the consortia websites.

PGC ADHD working group and the ADHD iPSYCH-SSI-Broad collaboration: https://www.med.unc.edu/pgc/results-and-downloads

ENIGMA and ENIGMA+CHARGE for ICV and hippocampus: http://enigma.ini.usc.edu/download-enigma-gwas-results/

## ACKNOWLEDGMENTS

Funding: This work was supported by the Netherlands Organization for Scientific Research (NWO), i.e. the NWO Brain & Cognition Excellence Program (grant 433-09-229) and the Vici Innovation Program (grant 016-130-669 to BF). Additional support was received from the European Community’s Seventh Framework Programme (FP7/2007 – 2013) under grant agreements n° 602805 (Aggressotype), n° 602450 (IMAGEMEND), and n° 278948 (TACTICS) as well as from the European Community’s Horizon 2020 Programme (H2020/2014 – 2020) under grant agreements n° 643051 (MiND) and n° 667302 (CoCA). The work was also supported by the Lundbeck Foundation (grant R102-A9118 and R155-2014-1724 to iPSYCH/A.D.B.), by a grant to the ENIGMA Consortium (grant number U54 EB020403) from the BD2K Initiative, a cross-NIH partnership, and by the ECNP Network ‘ADHD across the Lifespan’. SEM is supported by an Australian National Health and Medical Research Council fellowship (SRFB-1103623). This work was partly carried out on the Dutch national e-infrastructure with the support of SURF Foundation.

The Enhancing NeuroImaging Genetics through Meta-Analysis (ENIGMA) Consortium provided summary statistics of the consortium findings to this project. The original publication of those findings as well as the list of contributing samples and authors may be found on the ENIGMA website: http://enigma.ini.usc.edu. The Neurology Working Group of the Cohorts for Heart and Aging Research in Genomic Epidemiology (CHARGE) Consortium also contributed with an independent set of summary statistics of consortium findings. The contributing cohorts and authors may be found on previous publications of the CHARGE consortium as listed on the website: http://www.chargeconsortium.com/. The ADHD working group of the Psychiatric Genomics Consortium (PGC) and the iPSYCH-SSI-Broad collaboration ADHD Working Group contributed with an independent set of summary of consortium findings. The data and a complete list of contributing samples and people can be obtained from the PGC download website (https://www.med.unc.edu/pgc/results-and-downloads).

## AUTHOR CONTRIBUTIONS

Study conception and supervision: M.K., A.A.-V., B.F. Obtained funding: B.F., A.D.B. Provided samples and/or data: D.D, D.P.H., H.H.A., M.M., S.E.M. Conducted analyses: M.K. Contributed to analyses and data interpretation: R.K.W., J.L.S. Writing group: M.K., S.V.F., A.A.-V., B.F., A.D.B., J.B., N.R.M., R.S., E.S.-B., P.M.T. All authors reviewed and approved the final version of this manuscript.

